# Waking experience modulates sleep need in mice

**DOI:** 10.1101/2020.07.25.219642

**Authors:** Linus Milinski, Simon P. Fisher, Nanyi Cui, Laura E. McKillop, Cristina Blanco Duque, Gauri Ang, Tomoko Yamagata, David M. Bannerman, Vladyslav V. Vyazovskiy

## Abstract

Homeostatic regulation of sleep is reflected in the maintenance of a daily balance between sleep and wake. Although numerous internal and external factors can influence sleep, it is unclear whether and to what extent the process that keeps track of time spent awake is determined by the content of the waking experience. We hypothesised that alterations in environmental conditions may elicit different types of wakefulness, which will in turn influence both the capacity to sustain continuous wakefulness as well the rates of accumulating sleep pressure. To address this, we performed two experiments, where we compared wakefulness dominated by novel object exploration with either (i) the effects of voluntary wheel running (Experiment 1) or (ii) performance in a simple touchscreen task (Experiment 2). We find that voluntary wheel running results in longer wake episodes, as compared with exploratory behaviour; yet it does not lead to higher levels of EEG slow wave activity (SWA) during subsequent sleep. On the other hand, engagement in a touchscreen task, motivated by a food reward, results in lower SWA during subsequent sleep, as compared to exploratory wakefulness, even though the total duration of wakefulness was similar. Overall, our study suggests that sleep-wake behaviour is highly flexible within an individual, and that the homeostatic process that keeps track of time spent awake is sensitive to the nature of the waking experience. We therefore conclude that sleep dynamics are determined, to a large degree, by the interaction between the organism and the environment.

## Introduction

The duration, timing and intensity of sleep and wakefulness are under strict homeostatic control, and time spent awake is considered a key determinant of sleep pressure (Borbély, 1982). In addition, most animals partition the 24 hour day into consolidated periods of wakefulness and sleep with respect to environmental conditions, such as the light-dark cycle or food availability (Northeast *et al.*, 2019; Tam, Bannerman and Peirson, 2020). Typically, laboratory mice wake up near dark onset (ZT12) and remain awake for a variable duration before entering their first sleep bout. Evidence suggests that the duration of spontaneous wakefulness can be influences by essential homeostatic needs, motivated behaviours and environmental factors (Sotelo *et al.*, 2020). For example, providing access to running wheels results in an increased capacity to sustain continuous wakefulness (Vyazovskiy and Tobler, 2012; Fisher *et al.*, 2016). Furthermore, mice have also been shown to have longer waking bouts when food deprived (Northeast *et al.*, 2019).

Although it has been appreciated that environmental factors play an important role in understanding behaviour in general (Gomez-Marin and Ghazanfar, 2019) and sleep in particular (Ungurean *et al.*, 2020), surprisingly little is known about the effect of different behaviours elicited by environmental demands on sleep homeostasis. It is well established that sleep following prolonged wakefulness is characterised by elevated levels of electroencephalogram (EEG) slow-wave activity (EEG spectral power between 0.5-4 Hz), relative to the duration of preceding waking (Huber *et al.*, 2004; Vyazovskiy *et al.*, 2006). However, in addition to time spent awake evidence suggests that the specific nature of wake behaviours or activities may also influence subsequent sleep. For example, it was established that cortical SWA increases locally in the regions that were stimulated during preceding wakefulness (Vyazovskiy, Borbély and Tobler, 2000; Huber *et al.*, 2004), suggesting that sleep is regulated in a local, activity-dependent manner (Krueger *et al.*, 2019). Localised sleep-like activity has also been identified during wakefulness, particularly after prolonged wakefulness or extended training (Vyazovskiy *et al.*, 2011; Hung *et al.*, 2013; Bernardi *et al.*, 2015). Consistent with the detrimental effects of sleep pressure on cognitive performance (Krause *et al.*, 2017), these localised changes in cortical activity correlate with performance errors (Vyazovskiy *et al.*, 2011; Nir *et al.*, 2017), suggesting that the expression of local sleep is incompatible with sustained waking performance. Yet a question remains: which aspects of waking behaviour lead to the accumulation of sleep pressure?

Previous studies that investigated the relationship between waking activities and sleep mechanisms often utilised tasks that required precise motor coordination (Vyazovskiy *et al.*, 2011; Ramanathan, Gulati and Ganguly, 2015; Nagai *et al.*, 2017) or were designed to trigger neuronal plasticity mechanisms (Hanlon *et al.*, 2009; Hung *et al.*, 2013; Vyazovskiy and Harris, 2013; Yang *et al.*, 2014). Therefore, the observed changes in sleep/wake dynamics may not be directly related to the amount of wakefulness per se, and could instead arise as a result of the cognitive and/or attentional load of the tasks and their related neural activity (Tononi and Cirelli, 2006; Vyazovskiy and Harris, 2013). It is possible that engaging in less cognitively demanding behaviours while awake may have a very different effect on the accumulation of sleep pressure. This hypothesis is supported by evidence that sufficient training in a task may allow performance to become stereotypical and to require sustained activity only within restricted brain networks (Bueichekú *et al.*, 2016). Execution of stereotypical tasks can become independent of the primary motor cortex (Kawai *et al.*, 2015), or, as has been shown in song-birds, the forebrain (Bottjer, Miesner and Arnold, 1984). Thus, it is possible that a well-trained task involving the execution of a stereotypical motor-sequence rather than constantly adapting behaviour may be sustained even during localised changes in cortical activity or ‘local sleep’ in some cortical areas. Since evidence suggests that local cortical neuronal activities represent an important factor in governing global sleep homeostasis (Thomas *et al.*, 2020), engagement in well-trained behaviours may therefore result in an attenuated build-up of sleep need during waking. Consistent with this notion, it has been shown that stereotypic, repetitive wheel running is associated with a substantial prolongation of wake periods (Vyazovskiy *et al.*, 2006), yet this does not lead to an increase in cortical excitability, which was previously associated with increased sleep pressure (Huber *et al.*, 2013; Donlea, Pimentel and Miesenböck, 2014; Fisher *et al.*, 2016; Reichert, Pavón Arocas and Rihel, 2019). Although the neurophysiological mechanisms underlying the increased capacity to sustain spontaneous wakefulness are unclear, we previously proposed that wakefulness dominated by simple, stereotypic behaviours may be associated with reduced sleep need (Fisher et al., 2016).

Here we directly investigated the effects of two types of repetitive, stereotypic behaviour on sleep timing and EEG SWA during subsequent sleep. Specifically, we examined the effects of spontaneous running in a simple running wheel and voluntary performance in a well-learned, simple operant task involving food rewards. Both behaviours were compared to a condition in which animals were sleep deprived by providing novel objects to induce active, more demanding exploratory behaviour. We hypothesised that the nature of the waking behaviours has important influences on subsequent sleep, above and beyond its duration.

## Methods

### Experimental animals

Adult male C57BL/6J mice were used in this study (n=6-8 and n=5 in the first and the second experiment respectively, see below). The animals were individually housed in custom-made clear Plexiglas cages (20.3 × 32 × 35cm) with free access to a running wheel (RW). Cages were housed in ventilated, sound-attenuated Faraday chambers (Campden Instruments, Loughborough, UK, two cages per chamber) under a standard 12:12 h light-dark cycle (lights on 0900, ZT0, light levels ~120-180 lux). In experiment 1 food and water were available *ad libitum*, while in experiment 2 moderate food restriction (animals were kept at 85-90% of their average free feeding weight) was utilised to motivate performance. Room temperature and relative humidity were maintained at 22 ± 1°C and 50 ± 20%, respectively. Mice were habituated to both the cage and the cables connected to the cranial implant for a minimum of four days prior to recording. All procedures conformed to the Animal (Scientific Procedures) Act 1986 and were performed under a UK Home Office Project Licence in accordance with institutional guidelines.

### Surgical Procedures and Electrode Configuration

Surgical procedures were carried out using aseptic techniques under isoflurane anaesthesia (3-5% induction, 1-2% maintenance). During surgery, animals were head fixed using a stereotaxic frame (David Kopf Instruments, CA, USA) and liquid gel (Viscotears, Alcon Laboratories Ltd, UK) was applied to protect the eyes. Metacam (1-2 mg/kg, s.c., Boehringer Ingelheim Ltd, UK) was administered preoperatively. EEG screws were placed in the frontal (motor area, M1, AP +2mm, ML +2mm) and occipital (visual area, V1, AP −3.5-4mm, ML +2.5mm) cortical regions. A reference screw electrode was placed above the cerebellum. EEG screws were soldered (prior to implantation) to custom-made headmounts (Pinnacle Technology Inc. Lawrence, USA) and all screws and wires were secured to the skull using dental acrylic. Two single stranded, stainless steel wires were inserted either side of the nuchal muscle to record electromyography (EMG). At the end of the surgery animals were administered saline (0.1ml/20g body weight, s.c.). Thermal support was provided throughout the surgery and during recovery afterwards. Metacam (1-2 mg/kg) was orally administered for at least three days after surgery. A minimum two-week recovery period was permitted prior to cabling the animals.

### Signal processing and analysis

Data acquisition was performed using the Multichannel Neurophysiology Recording System (TDT, Alachua FL, USA). Cortical EEG was recorded from frontal and occipital derivations. EEG/EMG data were filtered between 0.1-100 Hz, amplified (PZ5 NeuroDigitizer pre-amplifier, TDT Alachua FL, USA) and stored on a local computer at a sampling rate of 257Hz (experiment 1) or 305Hz (experiment 2), and subsequently resampled offline at a sampling rate of 256 Hz. Signal conversion was performed using custom-written Matlab (The MathWorks Inc, Natick, Massachusetts, USA) scripts. Signals were then transformed into European Data Format (EDF). For each recording, EEG power spectra were computed by a Fast Fourier Transform (FFT) routine for 4-s epochs (fast Fourier transform routine, Hanning window), with a 0.25 Hz resolution (SleepSign Kissei Comtec Co, Nagano, Japan). All spectral analysis was based on signals acquired from the frontal EEG derivation, with the exception of one individual mouse in experiment 1, in which the occipital derivation was used, because the frontal signal was lost for technical reasons.

### Statistical analysis

All data are expressed as mean±standard error of the mean. Statistical analysis was performed using the Matlab (The MathWorks Inc, Natick, Massachusetts, USA) Statistics and Machine Learning Toolbox and custom written Matlab scripts. Wilcoxon rank sum tests were applied for comparing effects between any two experimental conditions without an additional time domain. Datasets with an additional time domain were tested with repeated measures ANOVA and Tukey post hoc tests for specific comparisons between groups. The level for statistical significance was set to p<0.05.

### Scoring and analysis of vigilance states

Vigilance states were scored offline through manual visual inspection of consecutive 4-s epochs (SleepSign, Kissei Comtec Co, Nagano, Japan). Two EEG channels (frontal and occipital) and EMG were displayed simultaneously to aid vigilance state scoring. Conventionally used criteria for vigilance state annotation were applied (Brown *et al.*, 2012; McKillop *et al.*, 2018). Vigilance states were classified as waking (low voltage, high frequency EEG with a high level or phasic EMG activity), NREM sleep (presence of slow waves, a signal of a high amplitude and low frequency particularly in the frontal EEG derivation) or REM sleep (low voltage, high frequency EEG dominated by theta frequency activity, with a low level of EMG activity, particularly in the occipital EEG derivation). Epochs contaminated by eating, drinking or gross movements resulting in artifacts in at least one of the two EEG derivations were excluded from analyses.

### Sleep deprivation

Animals were sleep deprived by manually introducing novel objects into their home cages at irregular intervals, as described previously (McKillop *et al.*, 2018). Novel objects included nesting material, small wooden blocks, and similar objects commonly used as environmental enrichment, and thus stimulated exploratory behaviour (exploratory wakefulness, EW). Each mouse was visually monitored throughout the period of sleep deprivation and the experimenter introduced objects whenever an animal showed signs of drowsiness, or stopped exploring. Sleep deprivation was undertaken for durations between approximately 1-6 hours depending on the experimental paradigm (see below).

### Experiment 1: The effects of voluntary wheel running on sleep

Running wheel (RW) activity is a widely used behavioural assay, utilising the well-described tendency of mice to spontaneously run in wheels when they are provided (Vyazovskiy *et al.*, 2006; Meijer and Robbers, 2014). Animals had free access to running wheels (Campden Instruments, Loughborough, UK, wheel diameter 14 cm, bars spaced 1.11 cm apart (inclusive of bars)) in their home cages for at least four weeks prior to the analysed dark period until ZT9 on each experimental day (Fig. 1), and were therefore well adapted to the wheels. The wheels were custom made for tethered animals, and tethering did not prevent the animals from running *ad libitum* (Fig. 2A). Each running wheel was fitted with a digital counter (Campden Instruments), which uses an infra-red (IR) emitter/receiver to detect each rung passing the IR beam as the wheel rotates. In our study RW activity was recorded with a high temporal resolution within the same system used to record electrophysiology signals, as reported previously (Fisher *et al.*, 2016). One full wheel revolution consisted of 38 individually detected rung counts, thus 10 counts per second corresponds to a speed of 10.11 centimeters/sec. The wheel counter output was a 5V TTL pulse (0V with no output) that triggered an edge detector in the TDT acquisition system, and in turn created a time stamp that was stored for each wheel count.

**Figure 1.**
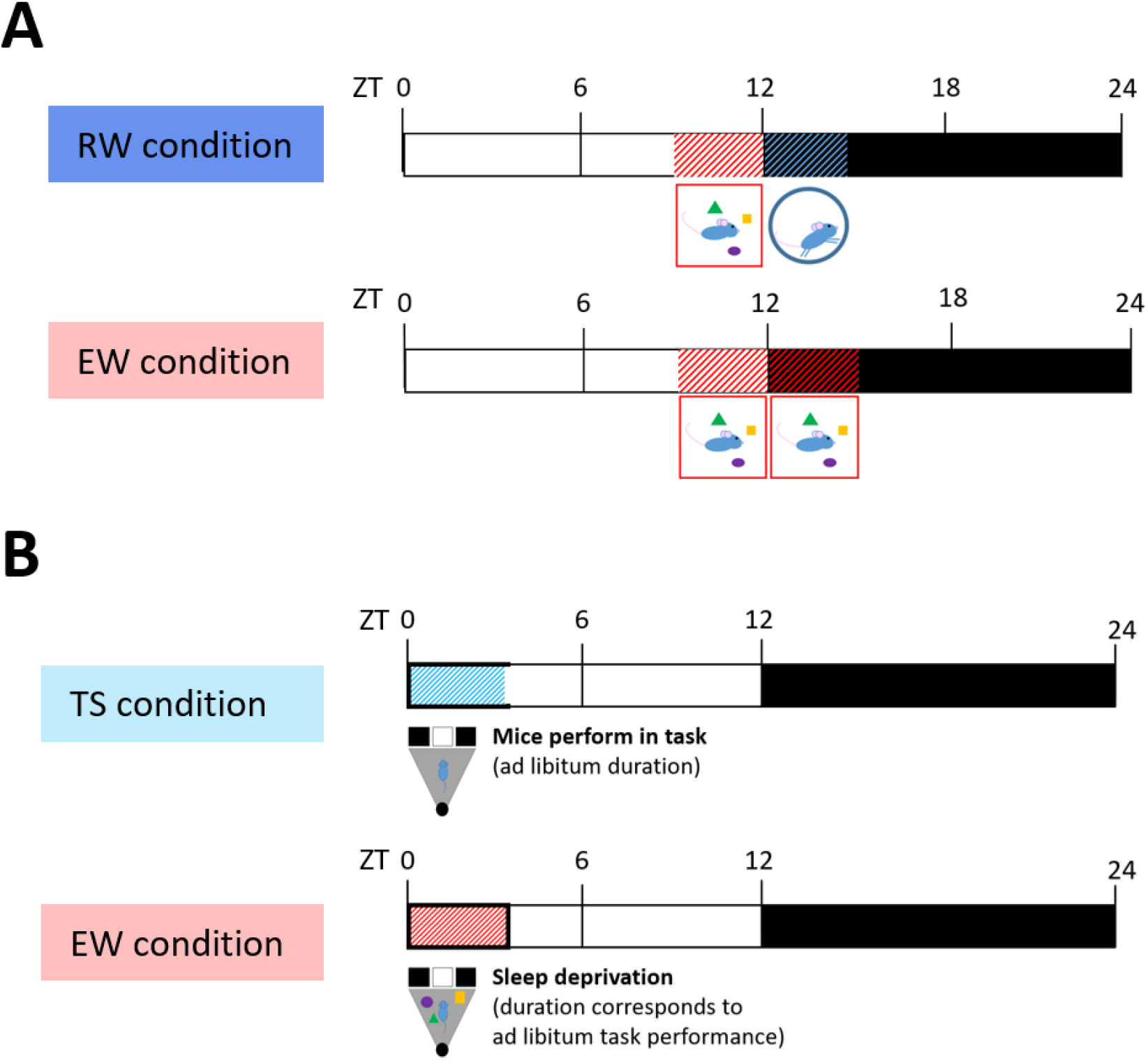
Schematic of experiment 1 and 2. **(A)** Experiment 1. Mice were tested in two experimental conditions. In both conditions, animals were first kept awake by providing novel objects for the last 3 hours of the light period (ZT9-12). Subsequently, in the RW condition, the mice were given unrestricted access to running wheels between ZT12-15, which they used voluntarily. In the EW condition mice were instead kept awake between ZT12-15 by providing novel objects, to elicit exploratory wakefulness. There was no access to running wheels in the EW condition. EEG recordings were acquired and analysed over the entire 24h period between ZT0-24 in either condition. **(B)** Experiment 2. In the TS condtion, mice were performing in the TS task for ad libitum duration starting from light onset. In the subsequent EW condition, mice were sleep deprived for the corresponding duration by providing novel objects to elicit exploratory wakefulness (EW). EEG recordings were acquired before and after task performance or sleep deprivation.

**Figure 2.**
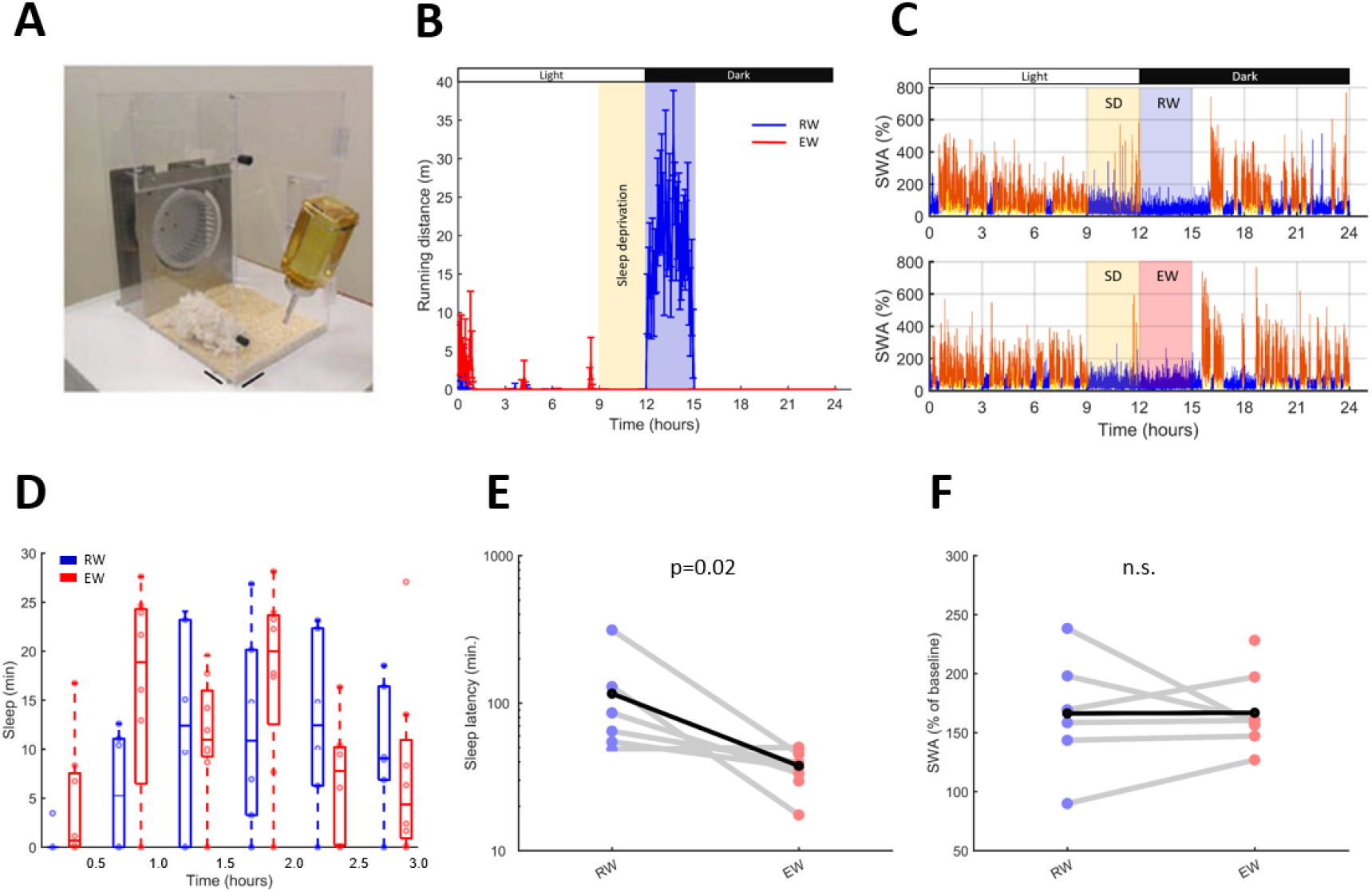
The effect of voluntary wheel running on sleep and SWA. **(A)** A photograph of a mouse home cage, fitted with a running wheel. EEG recordings were acquired continuously in the freely moving animals while in the home cage. **(B)** Time course of RW activity during the 24-h experiment, shown in 5-min bins. Note that the first 3 hours of the dark period (ZT12-15) are dominated by spontaneous wheel running in the RW-condition only. **(C)** Hypnogram of a representative mouse during the two experimental conditions. In the ‘RW’ condition (top) animals were sleep deprived for the last three hours of the light period and had access to the running wheel for the following 3 hours starting at dark onset. During the exploratory wakefulness (‘EW’) condition (bottom), animals were sleep deprived for the last 3 hours of the light period and then kept awake for further 3 hours starting at dark period, without access to the running wheel, but provided with novel objects to elicit exploratory behaviour. The plots depict color-coded EEG slow-wave activity (wakefulness: blue, NREM: red, REM sleep: yellow) with a 4s epoch resolution, shown as % of mean over the 24-h period. **(D)** Time course of total sleep during the first 3 h after the animals were left undisturbed (mean values, SEM). **(E)** Latency to the first consolidated sleep episode > 1 min. The dots indicate individual mice. **(F)** EEG SWA during the first 1-h interval from sleep onset. The dots indicate individual mice.

After habituation to the chamber, baseline recordings of undisturbed sleep and wakefulness spanning the 24 hours immediately before the experimental manipulation were performed. On the experimental day all animals were sleep deprived (SD) for the last 3 hours of the light phase without access to the running wheels and then subsequently kept awake for a further 3 hours from dark onset either by the introduction of novel objects to elicit predominantly exploratory behaviour (exploratory wakefulness, EW, n=8), or by providing access to the wheels (RW, n=6), which the mice used extensively (Fig. 1). At ZT15, 3 hours after dark onset, the novel objects were removed in the EW condition, and running wheels were blocked in the RW condition. All animals were then left undisturbed without wheel access for the rest of the dark period, and could sleep *ad libitum.*

This experiment was performed at the end of the light period in order for the animals to efficiently build up sleep pressure (by keeping them awake for the last 3h of their circadian rest phase) while ensuring voluntary wakefulness (either wheel running or exploratory behaviour) in the second half of the intervention, which took place at the beginning of their circadian active phase. Mice underwent both the RW and EW experimental conditions on separate days, though two mice were not recorded in the RW condition due to technical reasons (RW condition n=6 mice, EW condition n=8 mice).

### Experiment 2: The effects of touch screen performance on sleep

To test the effects of voluntary engagement in an operant, but not demanding behaviour on sleep we trained a separate group of adult male C57BL/6 mice (n=5) using the Bussey-Saksida Touch Screen system (Horner *et al.*, 2013; Heath, Bussey and Saksida, 2015). Our paradigm (called TS task) was designed specifically to allow for repetitive, stereotypical behavioural sequences to occur (Fig. 3A, Suppl. Video 1). Mice were first habituated to the training chamber and the milkshake (Yazoo strawberry milk drink) reward that was delivered to the reward tray at the rear of the chamber. Throughout the experiment, animals underwent food restriction to maintain their body weight at approximately 85-90% of their average body weight during *ad libitum* food access. Mice were fed daily after dark onset (between ZT12 and ZT15) so that feeding-related wakefulness (Northeast *et al.*, 2019) would coincide with their circadian active period.

**Figure 3.**
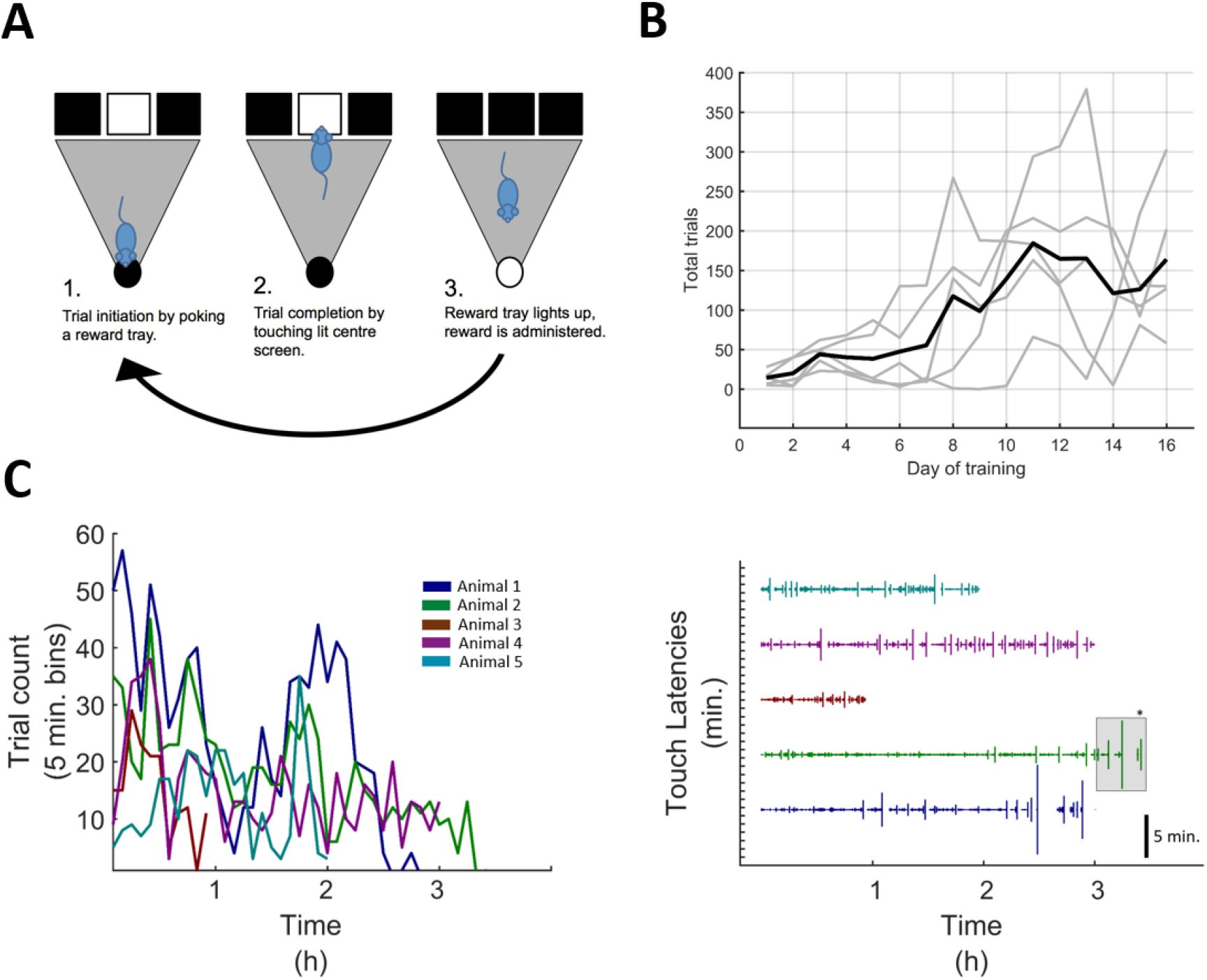
Voluntary wakefulness during an operant task. **(A)** Schematic of the operant task environment. A trial consists of nose-poking into a reward tray (1), followed by a single touch to an illuminated screen located in the centre of the opposite wall of the chamber (2). After completion of every three trials a food reward is administered into the reward tray. Collecting the food reward initiates the next trial (3). **(B)** Behavioural results of daily operant training, depicted as completed trials per session. **(C)** Behaviour during condition 1. Left: Behaviour shown as trial counts per 5 minute bin for each animal. Right: Behaviour depicted as latencies between touches to any screen or the food tray. Colours depict individual animals. Vertical lines depict latencies, × values of vertical lines depict onset of each measured latency (i.e. the last touch event). Stars for animal 4 (green) depict inactive time that was accounted for in the EW condition (i.e. the mouse was allowed to sleep after 3 hours).

In the TS task, a trial consisted of nose-poking into a reward tray, followed by a single touch to an illuminated screen located in the centre of the opposite wall of the chamber, which initiated the next trial. After completion of every trial, or, in later training stages, of every third trial, the milkshake reward (0.0035 ml,) was delivered to the reward tray. To encourage prolonged voluntary performance, there was no time limit for making a response in any part of the task. Note that trained animals still showed uniform response latencies across trials (Fig. 3C right panel). Daily training sessions were conducted over 2-4 weeks, typically lasting 30-45 min per day depending on how long it took for each mouse to complete at least 100 trials.

Mice then underwent two “experimental days” (Fig. 1B). On the first experimental day, the animals were disconnected and placed into the touch screen chamber at ZT0 (light onset) and allowed to perform the TS task continuously until they stopped engaging in the task for ten consecutive minutes (Fig. 3C), at which point they were transferred back into their home cage and reconnected to the recording setup. Pilot data from our lab suggested that animals may perform the task continuously for up to 7 hours or even longer (data not shown), and in the current study animals performed for an average of 2.5 hours. The experiment was conducted at the beginning of the light period as this is typically the start of the rest phase of mice and so performance in the task would be an effective form of sleep deprivation even for short task durations. Three to four days after the first experimental day the animals were again transferred into the touch screen chambers at light onset and kept awake for the same amount of time as they had previously performed the TS task, to match the environment and the time spent awake between the two conditions. Therefore, the duration of the voluntary performance in the TS task determined the duration of the sleep deprivation in the EW condition. This experimental design prevented us from counterbalancing the TS and EW experimental days. To control for effects of food restriction and feeding on sleep (Northeast *et al.*, 2019), each mouse in the EW condition was provided with the same type and volume of milkshake as in the previous TS task condition, administered at regular intervals throughout the sleep deprivation period. Continuous EEG and EMG recordings were conducted throughout the experiment, starting at least one day prior to the first experimental day and continuing until one day after the last experimental day. No electrophysiological recordings were obtained during engagement in either behavioural manipulation on the experimental days so as not to interfere with behavioural performance. Animals did not have access to running wheels during this experiment.

## Results

### Experiment 1: Voluntary wheel running increases the latency to sleep onset

In previous experiments, the effects of RW-activity on sleep were studied in mice with unrestricted access to wheels (Vyazovskiy and Tobler, 2012; Fisher *et al.*, 2016). Here, we compared wakefulness dominated by stereotypical wheel running to a more cognitively demanding exploratory behaviour (EW) in which animals were provided with novel objects to explore. Prior to the experiment until ZT9 on each experimental day, all animals had unrestricted access to running wheels (see Fig. 2A) and were well accustomed to running. As expected during the light period (typical rest phase of mice) before the experiment commenced, all animals showed negligible amounts of running (Fig. 2B), and a typical amount and distribution of sleep (representative mouse: Fig. 2C).

Mice were tested in two experimental conditions on separate days (Fig. 1A). On both experimental days all animals were kept awake using novel objects for three hours starting from ZT9 until the beginning of the dark period at ZT12 (Fig. 2B,C) while running wheels were blocked to prevent their use. At dark onset, in one condition, mice were sleep-deprived for an additional 3 hours, again by providing novel objects to elicit predominantly exploratory behaviour (exploratory wakefulness, EW). In the second condition mice were instead left undisturbed and given unrestricted access to the running wheel (RW), which all animals used extensively (on average 1552.6±251 wheel revolutions performed during the 3-h interval, which corresponded to 669.7±110.4 metres total distance covered, Fig. 2B). No novel objects were provided during this time, and the animals stayed awake spontaneously. Importantly, the total amount of sleep the animals obtained between ZT0-ZT9, prior to SD, was identical between the two conditions (RW: 6.12±0.2; EW: 6.12±0.3 hours, n.s.) as was the amount of sleep that occurred during 3-h SD between ZT9-12 (RW: 10.0±2.0; EW: 12.0±3.6 minutes, n.s.) or during the following 3 hour experimental manipulation between ZT12-ZT15 (RW: 4.5±2.1; EW: 1.5±0.8 minutes, n.s.). At ZT15, 3 hours after dark onset, all animals were then left undisturbed without wheel access for the rest of the dark period, and could sleep *ad libitum* (Fig. 2D). We found that the latency to sleep after the RW/EW procedure was significantly longer in the RW condition as compared to the EW condition (RW: 108.8±42.2; EW: 60.9±29.5 minutes, p=0.02, Wilcoxon rank sum test, Fig. 2E). However, the total amount of sleep during the first 1‐hr after sleep onset was similar between the conditions (EW: 37.6 ± 3.8 min, RW: 37.4 ± 4.8 min, n.s.), and the levels of NREM EEG SWA during this interval were nearly identical (EW: 166.9 ± 11.1%, RW: 166.0 ± 20.5% of mean between ZT0-ZT9, n.s., Fig. 2F). Thus, wakefulness dominated by wheel running delayed sleep onset by about an hour, without any effects on sleep pressure as measured by cortical EEG slow-wave activity.

### Experiment 2: Performance in a stereotypical touch screen task reduces the build-up of sleep pressure

Similar to Experiment 1, this paradigm was designed to address the effect of a simple, stereotypical behaviour (TS task) versus a more demanding exploratory wake behaviour on subsequent sleep (EW). However, in contrast to the RW-experiment, in which a certain amount of wakefulness (6-h) was enforced, this experiment was specifically designed to encourage voluntary wakefulness while animals performed a stereotypical task (TS task). The duration of the exploratory wakefulness condition was then time matched to the duration of the TS task.

To this end, mice (n=5) were trained in an operant touch screen (TS) task in a Bussey-Saksida Touch Screen chamber (Fig. 3A, Suppl. Video 1, see Methods for details). Prior to the experimental day animals were trained in the TS task for a minimum of 16 days. Animals showed a significant increase in the amount of trials over the course of training (F(15,60)=3.82, p<0.001, one way repeated measures ANOVA) and completed on average 164±23 trials on day 16 of training in comparison to 14±2 trials on day 1 (Fig. 3B).

After having completed at least 100 trials in a single session, animals performed the main experimental paradigm (TS) in which the mice were placed in the touch screen chamber at light onset (ZT0) and allowed to perform the task at will. Once the criteria for continuous, spontaneous disengagement from the task were met (see Methods, on average after 2.5 hours), the animals were transferred back to their home cages and undisturbed sleep recordings were performed until the end of the subsequent dark period (ZT24). Three to four days later, at light onset, the animals were again transferred into the TS operant chambers at ZT0 but instead of performing in the touchscreen task were sleep deprived (exploratory wakefulness, EW) for the same duration as they had each previously performed the TS task.

During unrestricted TS task performance, which lasted on average 2.5 hours (individual values: 1, 2, 3, 3 and 3.5 hours, n=5), the animals completed 156, 307, 544, 871 and 732 trials, respectively (Fig. 3C, left panel). As intended, the overall waking time (wake onset before the experimental manipulation to sleep onset afterwards) was nearly identical between the TS task and EW days (195±34min and 201±30min respectively, n.s., Fig. 4A, left panel) as was the total amount of sleep the animals obtained during the 12h dark period prior to the TS and EW days (TS 334.3±20.4 min., EW 279.9±31.2 min., n.s.) or during the preceding 12h light period (TS 444.7±32.5 min., EW 467.49±11.9 min, n.s., Wilcoxon rank sum test).

**Figure 4.**
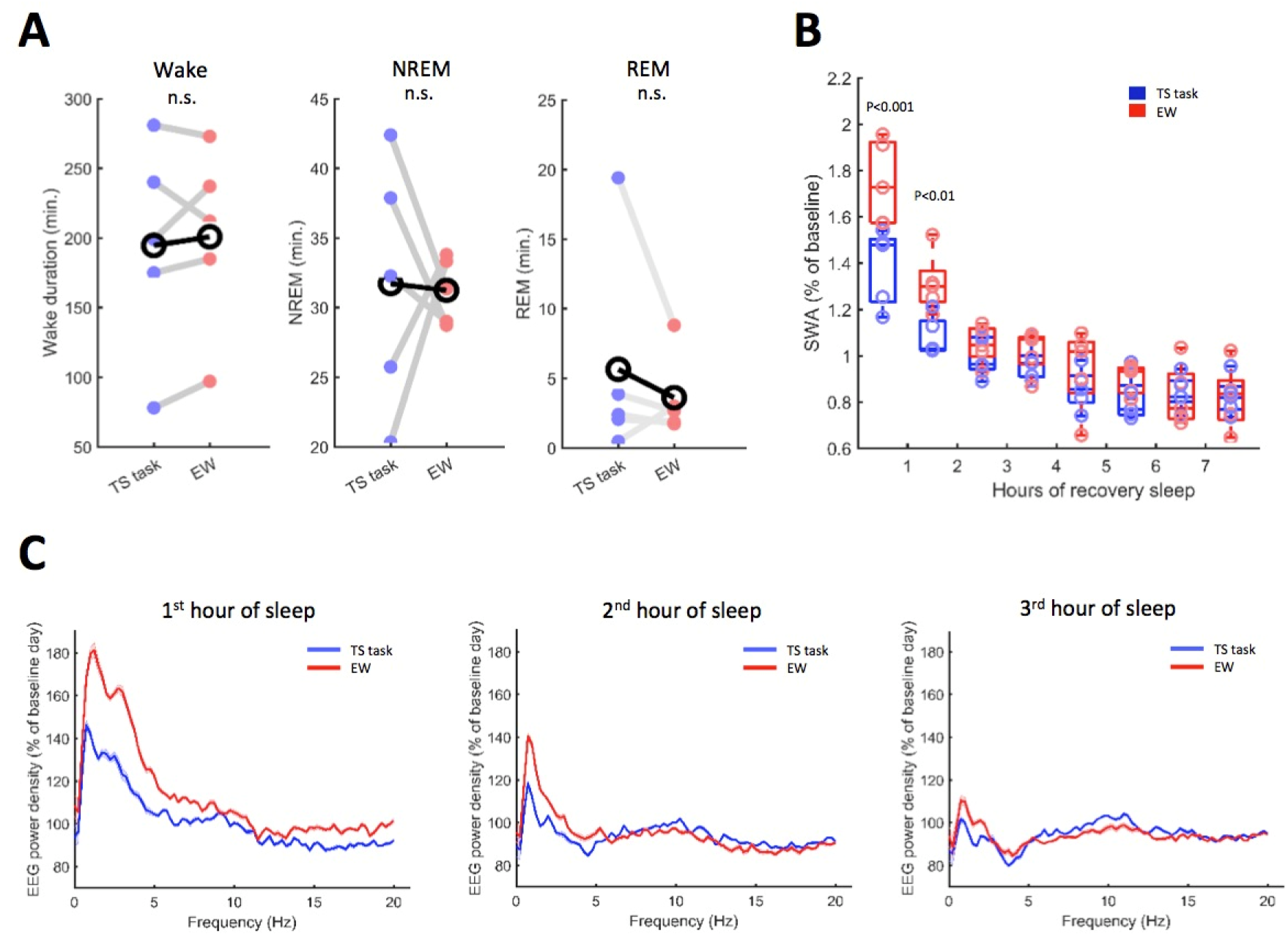
The effect of operant behaviour on sleep and SWA. **(A)** Sleep-wake pattern in the touchscreen task (TS task) and the corresponding exploratory wakefulness (EW) condition. Wake duration (left panel) describes duration of wakefulness starting from the end of consolidated sleep prior to the TS task or EW until sleep onset after the intervention. NREM and REM sleep duration (middle and right panel) depict durations within the first hour of recovery sleep after TS task (blue) or exploratory wakefulness. **(B)** Cortical EEG slow-wave activity (1-4 Hz EEG band) during recovery NREM sleep after the operant task (blue) and exploratory wakefulness (red). Points represent average SWA between individual animals shown in hourly bins, depicted as percentage of average SWA of preceding baseline day (interaction between hour and test condition: F(7,28)=11.824, p<0.01, repeated measures ANOVA). P-values in the graph correspond to post hoc Tukey test. **(C)** NREM EEG spectra of first, second and third hours of recovery sleep (from left to right) after the TS task (blue), and exploratory wakefulness (red). Shaded areas depict standard errors of the mean.

The amount of NREM and REM sleep during the first hour of sleep opportunity after the two conditions was not different between the ST and EW days (NREM: TS task 31.7±3.9min, EW 31.2±1.1min, n.s., REM: TS task 5.6±3.5min, EW 3.6±1.3min, n.s., Fig. 4A, middle and right panel). However, we observed that EEG slow wave activity was significantly higher during the initial two hours of recovery sleep in the EW condition as compared to TS task (interaction between factors ‘hour’ and ‘condition’ in two-way repeated measures ANOVA: F(7,28)=11.824, p<0.01; first hour EW 174.8±2.3% vs TS 138.6±2.1%, p<0.001, second hour EW 131.3±1.6% vs TS 108.3±1.1%, p<0.01, Tukey post hoc test, Fig. 4B). This effect was most prominent in frequencies within the slow-wave frequency range (Fig. 4C). The data suggest that voluntary engagement in a stereotypical operant task reduces the build-up of sleep pressure when compared to wake dominated by diverse exploratory behaviours.

## Discussion

Here we used two complementary experimental approaches to show that the nature of wake behaviour makes an important contribution towards the dynamics of sleep pressure, rather than this being determined only by wake duration. On the one hand, we show that wakefulness dominated by stereotypical wheel running behaviour results in a prolongation of spontaneous wakefulness without producing a higher level of EEG SWA during subsequent sleep. On the other hand, we demonstrate that voluntary engagement in an operant task that required repetitive, stereotypical behaviour only, reduces the levels of SWA during subsequent sleep, as compared to diverse exploratory behavior, despite the total wake duration being similar. We conclude that wakefulness associated with a low cognitive or attentional demand, may correspond to ‘waking with a lower cost’.

Although both types of repetitive, stereotypical behaviour investigated in this study were associated with a reduced accumulation of sleep pressure, their effects manifested in a different manner (i.e. prolonged wakefulness or decreased SWA during sleep). This was likely determined by the experimental design, including the time of day when the experiments were performed and the specific environmental manipulation(s) used. In the running wheel experiment the animals were kept awake either by providing novel objects while the running wheel was blocked or by providing unrestricted access to the wheel, which the animals used extensively. This kind of intervention was expected to be most efficient in the dark phase, which is the habitual wake period in mice. It allowed us to assess how long it would take for the animals to fall asleep spontaneously following exploratory behaviour or wheel running. We capitalised on the notion that wakefulness would be sustained until sleep pressure reached a certain upper threshold thereby initiating sleep. Consistent with our prediction, we found that the animals stayed awake significantly longer after wakefulness dominated by wheel running, as compared to exploratory wakefulness, yet the levels of SWA during subsequent sleep were identical between the conditions. On the contrary, in the touchscreen experiment, the animals stayed awake during the light phase by voluntarily performing a well-practiced, appetitively motivated, operant task (TS task). This experiment was performed during the light phase as this is the habitual sleep period of mice and so the total wake duration could be closely matched between the TS task condition and novel object EW condition. This allowed us to assess the levels of SWA during subsequent sleep after similar time spent awake but with very different waking experiences. As predicted, we found that SWA, which is an established measure of the homeostatic sleep pressure (Borbély, 1982; Guillaumin *et al.*, 2018), was significantly higher after EW as compared to TS performance.

It may be argued that the effects we observed in the TS condition (experiment 1) and, especially, in the running wheel condition (experiment 2) as compared to exploratory wakefulness may be due to increased physical activity of the mice. Previous research in human participants has shown that exercise can be sleep promoting, leading to increases in the proportion of NREM sleep and therefore exercise may be beneficial in conditions in which sleep is disturbed (Youngstedt, O’Connor and Dishman, 1997; Driver and Taylor, 2000; Kredlow *et al.*, 2015). Acute exercise before bedtime, specifically, has been shown to increase sleep amount (Myllymäki *et al.*, 2011; Aloulou *et al.*, 2019), but could also interfere with deep NREM sleep in the first hours of sleep (Aloulou *et al.*, 2019). We conclude that our findings of reduced sleep pressure after excessive wheel running or extended performance in the TS task as compared to exploratory wakefulness are in direct contrast to these previously reported sleep promoting effects of exercise. We also did not observe disruptions in NREM sleep specific to the early hours of sleep.

It is typically assumed that while the total sleep time differs greatly across the animal kingdom (Siegel, 2005), it is relatively stable within a species, suggesting that it is, at least to some extent, genetically determined (Andretic, Franken and Tafti, 2008). However, more recent evidence reveals a previously underappreciated flexibility with respect to the timing and duration of sleep (Lesku, 2012 *et al.*; Fisher *et al.*, 2016; Northeast *et al.*, 2019; Ungurean *et al.*, 2020), suggesting that extrinsic factors can also play an important role. Here we provide evidence to support the notion that the nature of waking behaviours affects the dynamics of sleep pressure, which in turn provides novel insights into the neurobiological substrates of sleep homeostasis (Borbély, 1982). Specifically, in this study we predictably altered the type of wake behaviour by manipulating environmental factors, which then affected the capacity to sustain continuous wakefulness or sleep intensity during subsequent sleep, depending on the time of day and task used.

Active wakefulness associated with higher neuronal activity (Thomas *et al.*, 2020) or EEG theta power (Vassalli and Franken, 2017) has previously been associated with increased homeostatic sleep pressure. The underlying neurobiological substrate of this association remains to be determined, yet its clear implication is that some behaviours may be associated with faster accumulation of sleep need. In support of this GluA1 AMPA receptor knockout mice have been shown to exhibit elevated SWA during sleep (Ang *et al.*, 2018) and increased EEG theta power in both the hippocampus and prefrontal cortex (and increased coherence of this theta activity across regions), which is tied specifically to elevated levels of exploratory attention in these mice (Bygrave *et al.*, 2019).

Conversely, other behavioural activities may lead to reduced or even negligible build-up of sleep need during waking. For example, wakefulness during food anticipatory activity in mice has been associated with reduced cortical indices of arousal and decreased EEG SWA (Northeast *et al.*, 2019). We propose that wakefulness dominated by repetitive or stereotypical motor sequences with lower attentional and/or cognitive demands may be the basis of a reduced build-up of sleep need. It has been previously suggested that, at least in some brain regions, neuronal activity during stereotypic running is functionally closer to sleep than to an awake state dominated by goal-directed purposeful behaviour (Fisher *et al.*, 2016). This notion is consistent with the finding that several animal species can reduce their sleep time dramatically when prolonged wakefulness is ecologically relevant or no longer optional, such as in some species of birds during migration (Rattenborg *et al.*, 2016; Rattenborg, 2017) or in marine mammals (Mukhametov, 1987; Lyamin, 1993; Lyamin *et al.*, 2008). It is likely, therefore, that substantial differences in the amount of sleep between species and at the individual level may be related to qualitative differences in waking behaviour, in addition to the need to satisfy fundamental biological drives such as reproduction (Lesku, 2012 *et al.*).

The lack of progress in establishing direct links between states of vigilance, overt behaviour and specific EEG activities has been acknowledged for some time. Major advancements over the last few decades have led to the appreciation that waking and sleep are highly dynamic states, which form a continuum, rather than representing distinct all-or-none conditions. While mixed states have been traditionally viewed as a maladaptive phenomenon that leads to impaired performance, we now propose that these states may have an adaptive value. For example, when sleep-like activities occur during wakefulness in brain areas not directly involved in task execution, essential behavioural performance could be maintained for extensive periods of time. This has previously been shown in great frigate birds who stay in the air for up to ten days while, intermittently, entering mostly unihemispheric sleep, correlated with stereotypical gliding flight but not manoeuvring (Rattenborg *et al.*, 2016; Rattenborg, 2017). Here we propose that the notion of maintaining certain wake behaviours during mixed states might not be specific to certain species but could be a wider phenomenon. Most likely candidates for waking behaviours compatible with mixed states are those that may not require coherent activation across large cortical networks, such as execution of stereotypical motor sequences, which have been shown to persist even with substantial cortical lesions (Bottjer, Miesner and Arnold, 1984; Kawai *et al.*, 2015).

In conclusion, we found that wakefulness consisting of the execution of a stereotypical, highly trained behaviour represents a state associated with a slow build-up of sleep pressure. Our data suggest a considerable flexibility in sleep-wake architecture and the dynamics of sleep homeostatic process, which are revealed when altered environmental conditions result in fundamental shifts of wake behaviour. Furthermore, we posit that wake duration and EEG slow-wave activity during sleep may be dissociated, suggesting a differential contribution of intrinsic and extrinsic factors in their respective control.

## Contributions

V.V.V. directed the study. V.V.V. and L.M. designed the experiments, analysed and interpreted the data and wrote the manuscript. L.M., S.P.F., N.C., L.E.M., C.B., T.Y. and G.A. installed experimental setups, performed the experiments and collected the data. D.M.B. provided advice on experimental design, data analysis and interpretation and contributed to writing the manuscript.

## Acknowledgements

Supported by: MRC NIRG (MR/L003635/1), BBSRC Industrial CASE grant (BB/K011847/1), FP7-PEOPLE-CIG (PCIG11-GA-2012-322050), John Fell OUP Research Fund Grant (131/032), Wellcome Trust Strategic Award (098461/Z/12/Z) and an Action on Hearing Loss studentship (S52_Bajo). We thank Drs Tim Bussey and Lisa Saksida for advice on touchscreen behavioural training in mice. We would like to thank Mathilde Guillaumin, Lukas Krone, Martin Kahn, Sibah Hasan, Yi-Ge Huang, Minas Salib, Christopher Thomas, Kathleen Reinhardt, Homero Esmeraldo and the whole Vyazovskiy group for support in surgeries, animal husbandry, conducting sleep deprivation and for many stimulating discussions.

## Notes

### Competing Interest Statement

The authors have declared no competing interest.

